# CISH, a novel intracellular immune checkpoint, in comparison and combination to existing and emerging cancer immune checkpoints

**DOI:** 10.1101/2025.04.26.650767

**Authors:** Florencia Cano, Alberto Bravo Blas, Mathilde Colombe, Chiara Cerrato, Ram Venegalla, Olivier Preham, Ellie Burns, Paige Mortimer, Nicholas Slipek, Matthew J Johnson, Beau R. Webber, Branden S. Moriarity, Emil Lou, Modassir Choudhry, Christopher A. Klebanoff, Tom Henley

**Author notes:** Co-corresponding Authors: Tom Henley,; Christopher A. Klebanoff.

## Abstract

Over the past decade, Immuno-Oncology has largely focused on blocking inhibitory surface receptors like PD-1 to enhance T cell anti-tumor activity. However, intracellular immune checkpoints such as CISH, which function independently of tumor-expressed ligands, offer powerful and previously untapped therapeutic potential. As a downstream regulator of TCR signaling, CISH controls T cell activation, expansion, and neoantigen reactivity. Though historically considered undruggable, recent advances in CRISPR engineering have enabled functional interrogation of these targets. We demonstrate that CISH deletion enhances T cell activation and anti-cancer functions more effectively than other emerging intracellular checkpoints. In CAR-T cells, CISH inactivation significantly increased sensitivity to tumor antigen, enabling robust recognition and killing even at low antigen levels, conditions that often lead to treatment failure with conventional T cell therapies, mirroring antigen escape scenarios seen in solid tumors. Our findings further validate CISH as a potent and druggable intracellular checkpoint capable of boosting anti-tumor T cell responses across diverse cancer types, independent of PD-L1 status. The underlying mechanisms of CISH inhibition may help explain the positive outcomes reported in recent clinical studies of this approach in solid tumor immunotherapy.

## INTRODUCTION

Cancer immunotherapies that boost the immune system’s ability to target tumors have transformed oncology, significantly extending survival in patients with advanced cancers. Among these, immune checkpoint inhibitors (ICIs) have become standard care for various malignancies^1, 2^. These therapies block inhibitory receptors like PD-1 and CTLA-4, which normally suppress T cell activity to maintain self-tolerance, with numerous inhibitors now commercially available^3, 4, 5, 6, 7, 8^. PD-1-targeting monoclonal antibodies are approved for several cancers including metastatic melanoma, non-small cell lung cancer (NSCLC), lymphoma and colorectal cancer (CRC)^5, 8^. However, their efficacy largely depends on PD-L1 expression in the tumor microenvironment, a factor that varies between patients and can decrease during treatment. Consequently, 30%–60% of patients fail to respond to PD-1/PD-L1 blockade^5, 9, 10^.

To improve clinical outcomes, a new class of intracellular immune checkpoints (ICs) are being explored both preclinically and clinically. Unlike conventional surface ICs that suppress T cell activity through ligand interactions, intracellular ICs act within the cell to inhibit TCR and cytokine signaling, offering potential for broad anti-tumor effects regardless of tumor type or biomarker status. We and others have identified CISH (cytokine-induced SH2 protein) as a key intracellular IC that is induced in CD8^+^ T cells following TCR stimulation, impairing anti-tumor function by targeting PLC-γ1 for proteasomal degradation^11, 12, 13^. It is markedly upregulated in tumor-infiltrating lymphocytes (TILs) compared to matched peripheral blood lymphocytes (PBLs)^11^. CISH depletion in patient-derived, neoantigen-selected TILs enhances proliferation, cytokine secretion, polyfunctionality, and tumor neoantigen recognition^11^. Notably, combining Cish knockout with PD-1 blockade in a preclinical murine model led to synergistic tumor regression^12^.

Historically, intracellular ICs have been difficult to target but advances in T cell gene-editing, such as CRISPR/Cas9, have opened new therapeutic possibilities. The anti-cancer potential of CISH depletion was recently tested in a first-in-human clinical trial for metastatic colorectal cancer (NCT04426669)^14^, using CRISPR-mediated CISH deletion. This study reported an exceptional complete response attributed to CISH knockout, underscoring CISH’s promise as a novel checkpoint addressing this unmet need and raising the prospect of small molecule inhibitors to broaden clinical access (Lou *et al*, *First-in-human clinical trial targeting the intracellular immune checkpoint CISH via CRISPR/Cas9-edited T cells in patients with metastatic colorectal cancers,* The Lancet Oncology*, April 29, 2025. Online ahead of publication*).

Building on these findings, we further investigated CISH and other emerging intracellular ICs using multiplexed gene-editing of primary human T cells, followed by functional assays in cancer antigen-specific *in vitro* systems.

## RESULTS

### Unlike PD-1, enhanced T cell effector and cytolytic functions by CISH inactivation are tumor cell surface ligand-independent

CISH differs from conventional immune checkpoints like PD-1, which rely on tumor ligand interactions, by acting intracellularly to suppress TCR signaling independently of target cell ligands^11^. Inhibiting CISH could therefore enhance anti-tumor responses across tumor types and biomarker statuses, crucial since many patients do not respond to PD-1/PD-L1 blockade^5, 9, 10^. Unlike CISH deletion, knockout of the *PDCD1* gene encoding PD-1 did not enhance T cell effector function or memory formation following αCD3 stimulation, suggesting that PD-1 inactivation alone does not confer benefit when TCR engagement occurs in the absence of its ligand, PD-L1. (**Fig. 1a–c, Suppl Fig. 1a,c**). To compare their functional impact, we evaluated CISH- and PD-1-deficient T cells in a KRAS^G12D^ cancer model using CD8^+^ T cells engineered to express a KRAS^G12D^-specific TCR via *TRAC* locus knock-in^15, 16^ (**Suppl Fig. 1c**). We selected the KRAS^G12D^ antigen because mutations in the KRAS oncogene are common and play a key role in the initiation and progression of many human cancers, making it a clinically relevant target for assessing T cell responses. The enhanced effector phenotype from CISH deletion persisted upon cognate antigen stimulation, either by peptide-pulsed H-1975 cells or direct TCR activation, even at low antigen levels (e.g.: 0.1 µg/mL), as shown by cytokine profiling (**Suppl Fig. 1d-e**). This enhanced effector phenotype was evident even in response to low levels of pulsed KRAS^G12D^ peptide (e.g.: 0.1 ug/ml) (**Suppl Fig. 1e**). Finally, we assessed cytolytic activity of KRAS^G12D^-TCR CD8^+^ cells against PD-L1/2-expressing H-1975 targets (**Suppl Fig. 1f**). CISH-deficient cells demonstrated superior antigen-specific killing compared to PD-1 KO cells, and this effect was independent of PD-L1/2 expression. (**Fig. 1d, Suppl Fig. 1f-i**).

**Figure 1.**
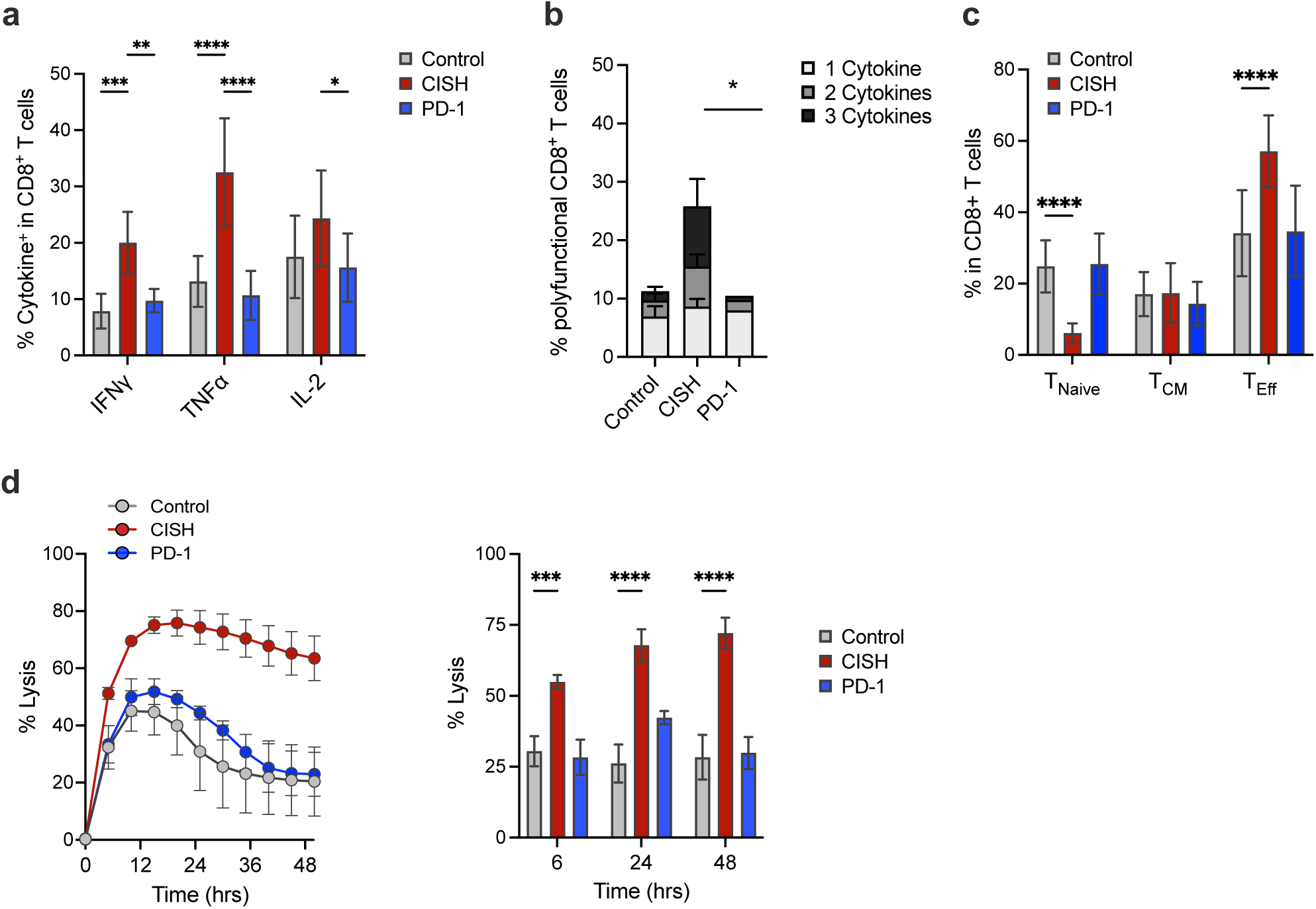
Unlike PD-1, enhanced T cell effector and cytolytic functions by CISH inactivation are tumor cell surface ligand-independent. **a**, Knockout of CISH significantly increases the magnitude of effector cytokine production measured by ICS after ⍺CD3 stimulation (1 ug/ml), whereas PD-1 knockout results in cytokine production similar to control cells. **b,** Contrary to CISH depleted CD8^+^ T cells, PD-1 knockout does not enhance cytokine polyfunctionality. **c**, The increase in T cell memory formation measured by flow cytometry after TCR stimulation, in particular effector memory cells, observed by knockout of CISH is not seen by PD-1 depletion. **d,** CISH depletion enhances KRAS^G12D^ antigen-specific cytolysis compared to control T cells, while loss of PD-1 shows no significant improvement in cytolysis. Kinetic impedance assay using xCELLigence enables real-time measurement of cytolysis of KRAS^G12D^ expressing H-1975 cells by edited KRAS^G12D^-TCR^+^ CD8^+^ T cells. (4:1 Effector: Target ratio). **a-d:** Statistical significance was determined by either One or Two-way ANOVA with multiple comparison test for repeated measures. Not shown = not significant, *P >0.05, **P>0.01, ***P>0.001, ****P>0.0001. All data are representative of three independent experiments. N=6. Data are mean ± SEM.

### CISH inactivation enhances T cell function more effectively than the inactivation of other prominent intracellular immune checkpoints

To compare the immune-enhancing effects of CISH knockout with other emerging intracellular immune checkpoints (**Fig. 2a)**, we developed a CRISPR/Cas9-based multiplex gene-editing strategy for primary human CD8^+^ T cells, followed by phenotypic analysis after TCR stimulation (**Suppl Fig. 2a**). We assessed whether CISH deletion increases T cell sensitivity to low-level TCR stimulation, mimicking low antigen expression in tumors, as previously observed in KRAS^G12D^-TCR CD8^+^ cells (**Suppl Fig. 1e**). CISH-deficient cells showed significantly elevated production of effector cytokines IFNγ, TNFα, and IL-2 (**Fig. 2b–f, Suppl Fig. 2e**), including higher IFNɣ expression levels and frequency (**Fig. 2b–d**), as well as increased cytokine polyfunctionality (**Fig. 2f**). These effects were most pronounced at low ⍺CD3 stimulation and sustained after 21 days in culture (**Fig. 2b–e, Suppl Fig. 2c–d**). CISH knockout also induced a stronger program of T cell activation (**Fig. 2h**), effector function (**Fig. 2b–f**), and effector memory T cell formation (**Fig. 2g**) than knockouts of other intracellular ICs, including RASA2, CBLB, SOCS1, REGNASE1, and HPK1, under low ⍺CD3 stimulation. Notably, unlike RASA2 and CBLB knockouts, PD-1 expression at low ⍺CD3 stimulation remained comparable between control and CISH-deleted cells (**Fig. 2i**), suggesting a distinct regulatory profile. These data demonstrate that CISH deletion more effectively enhances T cell activation upon TCR ligation than other intracellular immune checkpoints tested.

**Figure 2:**
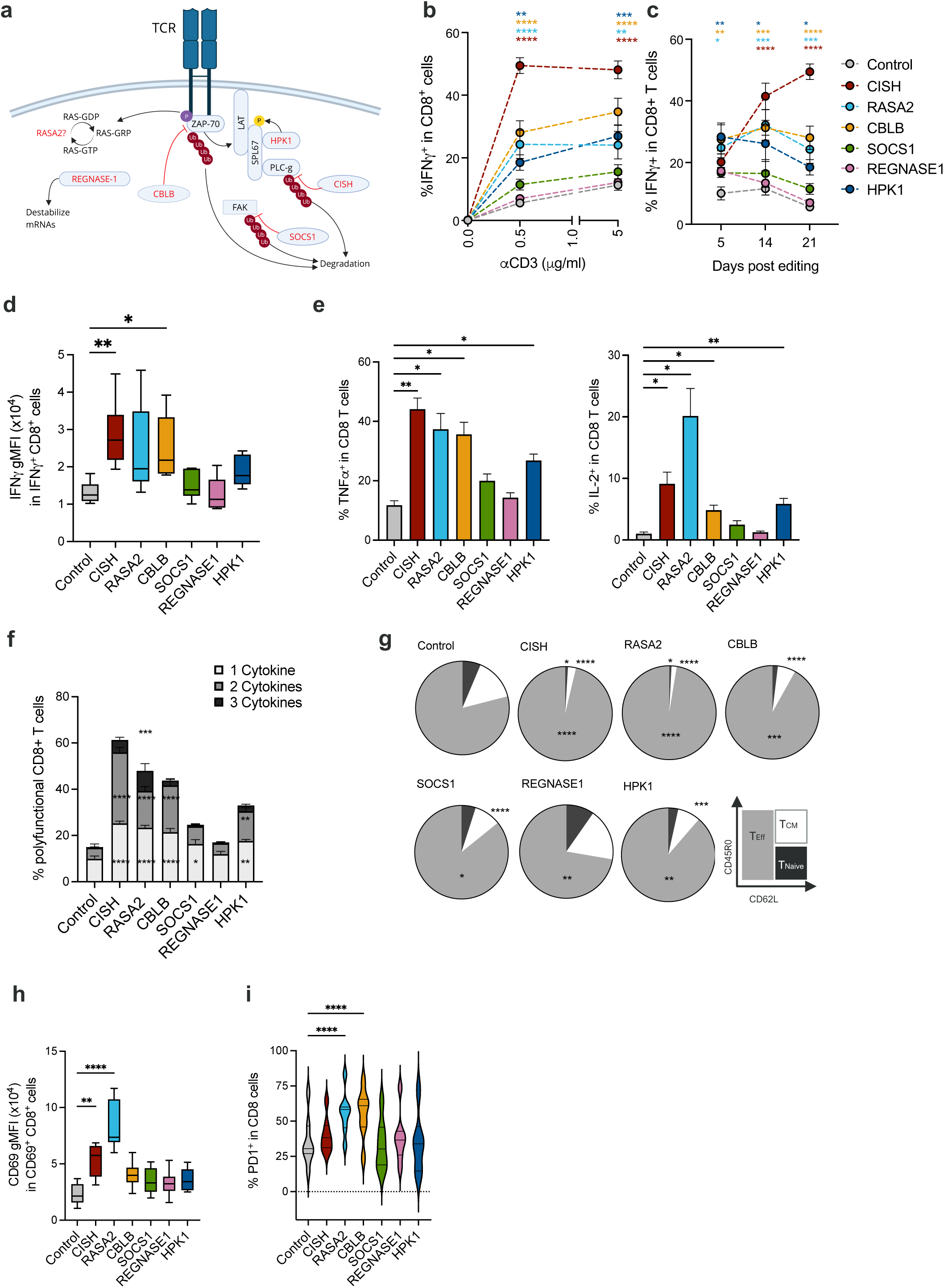
CISH inactivation enhances T cell function more effectively than the inactivation of other leading intracellular immune checkpoints. **a**, Intracellular immune checkpoint signaling in human T cells showing the unique and non-redundant roles of select checkpoint targets in the context of antigen receptor signaling. **b**, Effector Cytokines: IFN**γ** production at day 21 (D21) in CD8^+^ cells after increasing levels of ⍺CD3 stimulation, as measured by intracellular cytokine staining (ICS). Statistics shown vs. Control at each ⍺CD3 dose. **c,** Evaluation of IFN**γ** production in CD8^+^ cells upon 0.5 ug/ml ⍺CD3 stimulation, over time post editing. Statistics shown vs. Control at each time point. **d,** Intensity of intracellular IFN**γ** staining in stimulated immune checkpoint knockout (KO) cells (0.5 ug/ml ⍺CD3 stimulated cells, at day 21). **e,** Evaluation of TNF**α** and IL-2 production at day 21 upon 0.5 ug/ml ⍺CD3 stimulation. **f,** CISH KO in human CD8^+^ T cells increases the ratio of polyfunctional CD8^+^ T cells after TCR stimulation via ⍺CD3 stimulation (0.5 ug/ml, Day 21). Statistics shown vs. Control at each category. **g,** CISH KO in CD8^+^ T cells increases the proportion of cells with an effector memory phenotype at day 21 after editing upon 0.5 ug/ml ⍺CD3 stimulation. Statistical significance was determined by One-way ANOVA vs. Control in each population. **h,** CISH KO CD8^+^ T cells activation status is increased compared to control T cells upon 0.5 ug/ml ⍺CD3 stimulation (Day 21). **i,** Expression of inhibitory receptor PD-1 is similar between CISH KO and control T cells at low ⍺CD3 stimulation (0.5 ug/ml), Day 21). **b–i**: Statistical significance was determined by ANOVA vs. Control (unless otherwise stated); *P ≤ 0.05; **P ≤ 0.01; ***P ≤ 0.001; ****P ≤ 0.0001; not shown = not significant. All data is representative of at least three independent experiments, N=6. Error bars represent mean ± SEM.

### CISH functions as a non-redundant regulator of antigen-specific T cell cytolysis, and its inactivation can synergize with other intracellular immune checkpoints

The complexity of TCR signaling suggests that multiple non-overlapping pathways are governed by distinct intracellular ICs (**Fig. 2a**). To explore potential synergistic effects, we expanded our CRISPR multiplex editing platform in primary human T cells to test combinations of CISH depletion with other ICs (**Suppl Fig. 3a**). Using low ⍺CD3 stimulation (0.5 μg/ml), we observed an additive increase in cytokine production, notably IFNγ, TNFα, and IL-2, only when CISH was co-deleted with SOCS1, HPK1, or RASA2 (**Fig. 3a–b**), supporting non-redundant roles in regulating T cell effector functions. However, no additive effects were seen in the frequency of effector memory T cells or activation status with these combinations (**Suppl Fig. 3b–c**).

**Figure 3:**
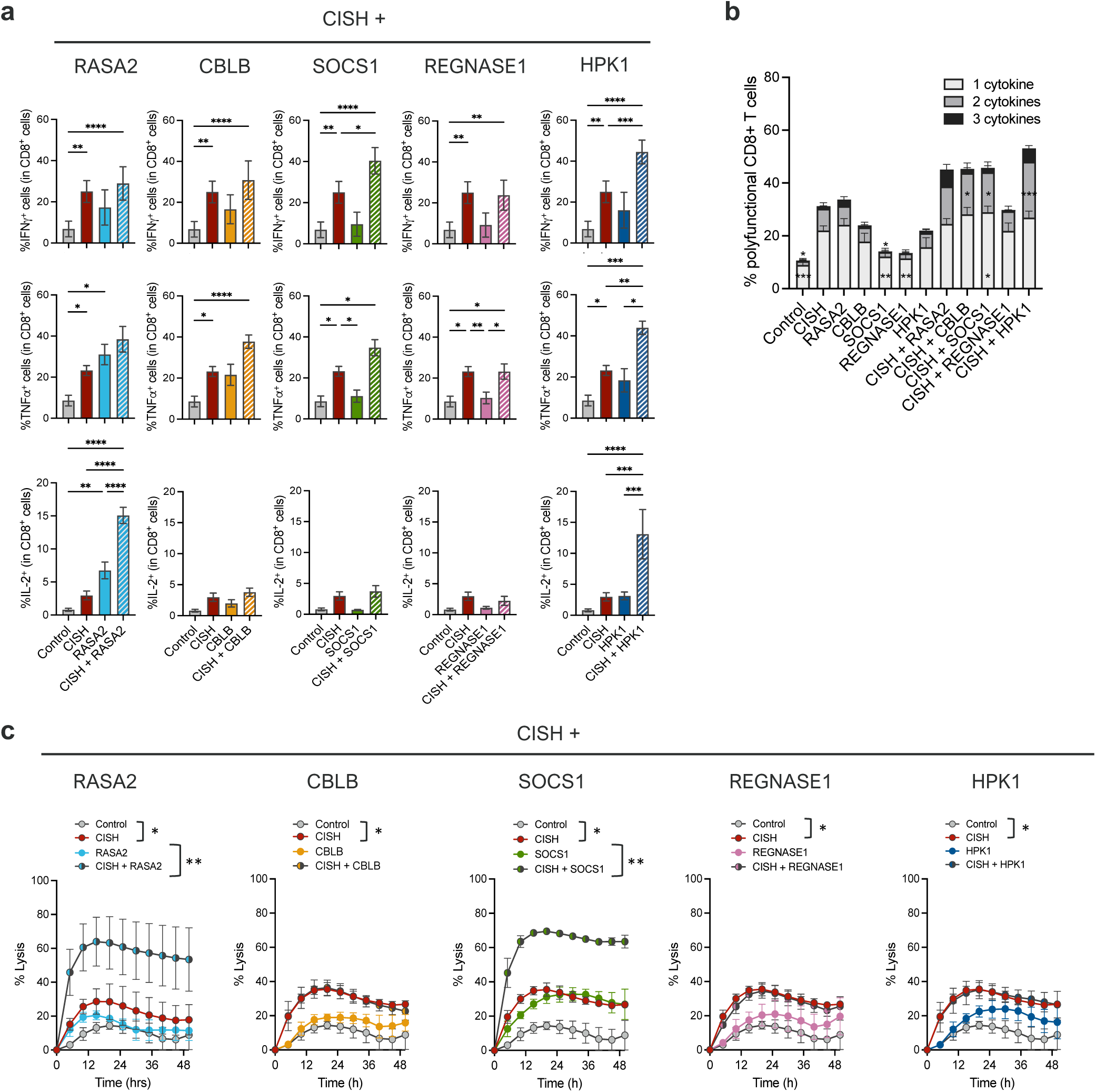
CISH functions as a non-redundant regulator of antigen-specific T cell cytolysis, and its inactivation can synergize with other intracellular immune checkpoints. **a**, Increase in T cell effector function when CISH is depleted in CD8^+^ cells in combination with other intracellular immune checkpoints. Effector cytokines production at day 21 post editing upon 0.5 ug/ml ⍺CD3 stimulation, measured by ICS. **b**, Polyfunctionality of CD8^+^ T cells after TCR stimulation via ⍺CD3 stimulation when CISH is depleted in combination with other intracellular immune checkpoints. **a, b**: Statistical significance was determined by One-way ANOVA vs. CISH KO. *P ≤ 0.05; **P ≤ 0.01; ***P ≤ 0.001; ****P ≤ 0.0001; not shown = not significant. All data is representative of at least three independent experiments, N=6. Error bars represent mean ± SEM. **c**, Antigen-specific killing in a model of KRAS^G12D^ solid tumor when CISH is depleted in combination with other intracellular immune checkpoints: Cytolysis of KRAS^G12D^ expressing H-1975 cells by edited KRAS^G12D^-TCR^+^ CD8^+^ T cells. (2:1 Effector: Target ratio). Statistical significance was determined by One-way ANOVA vs. CISH KO at 24hrs timepoint. *P ≤ 0.05; **P ≤ 0.01; ***P ≤ 0.001; ****P ≤ 0.0001; not shown = not significant. All data is representative of at least three independent experiments. Error bars represent mean ± SEM.

In the same KRAS^G12D^ antigen-specific solid tumor model, CISH deletion enhanced tumor cell killing by KRAS^G12D^-TCR^+^ T cells (**Fig. 3c**), and this effect was further potentiated by co-deletion with RASA2 or SOCS1, again underscoring the existence of distinct regulatory pathways. The synergy with SOCS1 is consistent with its role as a member of the same SOCS family as CISH. Notably, recent work by Schlabach *et al*.^17^ (KSQ-001EX) demonstrated that SOCS1 inactivation enhances anti-tumor activity in PD-1– refractory models and preclinical TIL studies, supporting the rationale for dual CISH/SOCS1 checkpoint inhibition in cancer therapy.

We also evaluated HPK1, a kinase under active clinical investigation as a small molecule checkpoint inhibitor^18,19,20^. Although co-deletion of CISH and HPK1 increased effector cytokine production (**Fig. 3a–b**), it did not enhance antigen-specific cytolysis (**Fig. 3c**), suggesting a partial functional synergy.

Additionally, we assessed PTPN1 and PTPN2, phosphatases targeted by novel small-molecule inhibitors currently in clinical trials (NCT04777994^21^, NCT04417465^22^). CISH deletion alone outperformed PTPN1, PTPN2, and combined PTPN1/2 knockouts in terms of T cell activation, cytokine production, polyfunctionality, and cytolytic activity (**Suppl Fig. 3a–d**). Co-deletion with CISH did not yield additive benefits in cytolysis, indicating that CISH operates through distinct, dominant pathways in regulating anti-tumor T cell function.

### CISH inactivation enhances CD19-specific CAR T cell cytotoxicity against low-antigen expressing cancer cells

Adoptive cell therapies (ACT) using tumor-specific T cells have shown strong clinical efficacy in certain hematological malignancies^23, 24^, but success in solid tumors remains limited^25, 26^. To assess the translational relevance of CISH deletion, we extended our prior work with KRAS^G12D^-TCR T cells to a CD19-CAR T cell model targeting hematologic cancers. Primary human CD8^+^ T cells were retrovirally transduced with a second-generation CD19-specific CAR incorporating a CD28 costimulatory domain^27, 28^ (**Suppl Fig. 5b**). These CD19-CAR^+^ T cells were then co-cultured with NALM6 B-cell leukemia cells engineered to express varying levels of CD19 (**Suppl Fig. 5a**), and cancer cell killing was measured by luciferase activity. CISH knockout significantly enhanced tumor cell lysis (**Fig. 4a–b, Suppl Fig. 5c**) and antigen-specific IFNɣ production (**Fig. 4c**), including at low CD19 expression levels, recapitulating the enhanced sensitivity observed previously (**Fig. 2b, Suppl Fig. 1e**). Similar results were obtained in a non-B cell line (H-1975) expressing CD19, confirming that CISH inactivation broadly boosts antigen-specific CAR-T cytotoxicity (**Suppl Fig. 5d–e**).

**Figure 4:**
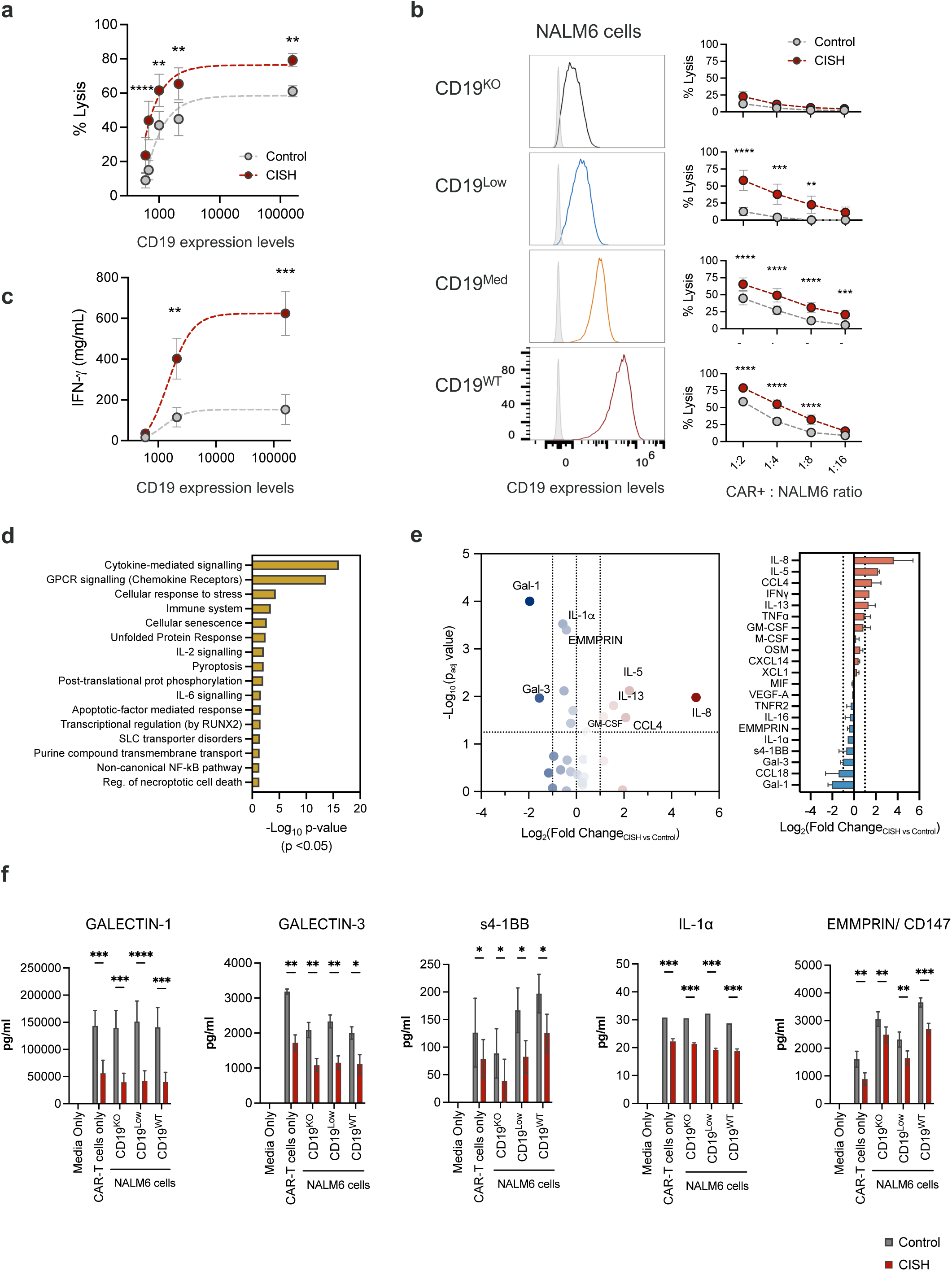
CISH inactivation enhances CD19-specific CAR T cell cytotoxicity against low-antigen expressing cancer cells. **a**, NALM6 target cell killing by CD8^+^ CD19-CAR^+^ T cells in the presence (Control) or absence of CISH (CISH) at effector: target ratio 1:2. **b**, Left, CD19 expression (colored) on engineered NALM6 target cells compared with unstained cells (grey). Right, CD19-CAR^+^ T cell killing of NALM6 cells expressing varying levels of CD19. **c**, Effector cytokine (IFN**γ**) levels in CD19 stimulated CD19-CAR^+^ CD8^+^ cells, as measured by ELISA. **a–c**: Statistical significance was determined by 2-way ANOVA vs. Control: Not shown = not significant, *P ≤ 0.05; **P ≤ 0.01; ***P ≤ 0.001; ****P ≤ 0.0001. All data is representative of at least three independent experiments. N=10 donors. Error bars represent mean ± SEM. **d–f**, Cytokine profile of CISH depleted CD19-CAR^+^ T cells after overnight co-culture with CD19 expressing NALM6 cells (as measured by nELISA). **d**, Pathway analysis by Reactome.org. Graph shows significantly regulated pathways; p≤0.05 (x-axis: Pathway Hierarchy/ Go Biological Process; y-axis; -Log_10_(p-value)). **e,** Volcano plot (Log_2_FC vs. -Log_10_(p-value); Left panel) and bar graph (Log_2_FC ≥ 1.2); Right panel) of differentially regulated secreted factors in CISH KO vs. Control CD19-CAR^+^ CD8^+^ cells after overnight co-culture with CD19^WT^ NALM6 cells. **f,** Significantly downregulated factors in CISH depleted CD19 stimulated CD19-CAR^+^ CD8^+^ cells. **d-f**: Statistical significance was determined by either multiple t-test or 2-way ANOVA vs. Control in each treatment. Not shown = not significant, *P ≤ 0.05; **P ≤ 0.01; ***P ≤ 0.001; ****P ≤ 0.0001. N=3 donors. Data are mean ± SD.

To further characterize this phenotype, we performed a nELISA cytokine screen (Nomic Bio) after overnight co-culture of CD19-CAR^+^ T cells with CD19^+^ NALM6 cells (**Fig. 4d–f**). CISH-deficient CAR-T cells exhibited elevated production of IFNɣ, TNFα, IL-5, CCL4, and GM-CSF, cytokines associated with T cell activation, tumor infiltration, and immune recruitment (**Fig. 4d–e**). Interestingly, CISH deletion also led to reduced secretion of immunosuppressive or tumor-promoting factors (**Fig. 4e–f**), including: Galectins 1 and 3, which negatively regulate T cells and induce apoptosis^29, 30^; Soluble 4-1BB (sCD137), known to attenuate co-stimulatory signaling^31, 32^; IL-1α, a DAMP molecule shown to inhibit CD8^+^ memory formation and IFNɣ production^33,34^; and EMMPRIN (CD147), a tumor-associated glycoprotein that promotes immunosuppressive cytokine release and correlates with T cell exhaustion^35, 36, 37^.

Collectively, these findings suggest that CISH inactivation enhances CAR-T cell function in part by increasing antigen sensitivity and effector function while reducing exposure to immunosuppressive factors. This makes CISH targeting a compelling strategy to improve T cell performance, particularly for tumors with low antigen density or immunosuppressive microenvironments.

## DISCUSSION

While single and combination therapies targeting cell surface immune checkpoints such as PD-1 and CTLA-4 have produced notable clinical responses, a significant proportion of patients remain unresponsive or ultimately relapse. In this study, we demonstrate that genetic ablation of CISH enhances antigen sensitivity and sustains effector function in engineered human T cells. Unlike conventional surface checkpoints, CISH belongs to a novel class of intracellular regulators that act downstream of the TCR and function independently of ligand engagement. This ligand-independent mechanism offers a strategic advantage, as targeting CISH may overcome multiple upstream inhibitory signals simultaneously, regardless of tumor type or biomarker expression. These findings support the therapeutic promise of CISH inhibition as a broadly applicable and potentially superior approach to current immune checkpoint blockade strategies.

Several emerging intracellular immune checkpoints have been proposed as promising targets to enhance T cell-based immunotherapies^17, 38, 39, 40, 41, 42^. Notably, Carnevale *et al*. demonstrated that RASA2 depletion in NY-ESO-1-specific TCR-T cells, as well as in CD19- and EphA2-targeting CAR-T cells, significantly improved tumor control and survival in both hematologic and solid tumor models^38^. Similarly, Wei *et al*. identified REGNASE1 as being highly expressed in tumor-infiltrating lymphocytes (TILs) and showed that its deletion enhanced adoptive cell therapies (ACTs) by promoting the development of long-lived effector T cell phenotypes^41^. While these regulators act at distinct nodes of the T cell activation network, their functional roles partially overlap or converge on shared signaling cascades, such as MAPK, JAK-STAT, and NF-κB, resulting in redundancy in maintaining immune homeostasis and preventing overactivation. In direct comparison, CISH ablation induced a more robust program of T cell activation, marked by enhanced antigen-specific cytolysis, effector cytokine production, and effector memory T cell formation. These improvements were consistently superior to those observed with other intracellular ICs, including RASA2, CBLB, SOCS1, REGNASE1, HPK1, and PTPN1/2, particularly under conditions of low antigen expression, a common feature of tumor escape. Additionally, our findings demonstrate that the concurrent inactivation of CISH and select ICs (RASA2 and SOCS1) can act in a non-redundant synergistic manner to significantly further improve the tumor-specific cytolytic potential of T cells, offering the compelling prospect of a dual-therapeutic approach. Importantly, CISH appears to exhibit a particularly broad suppressive role, integrating both cytokine and TCR signaling, and may offer a more comprehensive strategy to enhance T cell efficacy compared to single-pathway regulators.

The finding that CISH ablation enhances T cell effector and cytolytic functions downstream of distinct antigen receptors, including both chimeric antigen receptors and T cell receptors (e.g.: KRAS^G12D^-specific), further supports the receptor-independent nature of CISH biology. This highlights the broad therapeutic potential of CISH inhibition to enhance the anti-tumor activity of diverse T cell modalities, whether engineered CAR-T cells, patient-derived tumor-infiltrating lymphocytes, or endogenous peripheral tumor antigen-specific T cells. Most importantly, CISH deletion significantly increased T cell sensitivity to antigen, a benefit observed in *in vitro* models using CAR- and TCR-engineered T cells. In these contexts, CISH-deficient T cells were able to recognize and kill tumor cells expressing even very low levels of antigen, conditions that closely mirror tumor antigen escape, a key mechanism by which cancers evade conventional T cell therapies.

CISH ablation in CD19-CAR^+^ T cells also showed significantly reduced levels of secreted Galectins 1 and 3, even in the absence of CD19 stimulation, perhaps due to impact on cytokine receptor signaling. Galectins (Gal) are a family of mammalian β-Galactoside-binding proteins that play a key role in T cell homeostasis in the TME by acting as negative extracellular checkpoints of T cell function, inducing lymphocyte deactivation, and promoting T cell death^29, 30^. Indeed, both Gal-1 and Gal-3 inhibition significantly reduces tumor growth, increases activated CD8^+^ infiltration into the TME, and increases IFNɣ secretion from CD8^+^ TILs improving their cytotoxicity^43, 44, 45, 46^. Increasing attention has been paid on Gal-1 and Gal-3 inhibitors which have shown enormous potential in tumor therapy with Gal-1 inhibition boosting anti-PD1 therapy responses^30, 43^.

The most compelling evidence supporting the therapeutic potential of targeting CISH comes from the first-in-human clinical trial employing CRISPR-mediated knockout of *CISH* in T cells administered to patients with metastatic colorectal cancer (NCT04426669^14^). In this study, CISH-knockout Tumor-Infiltrating Lymphocytes were used as a model system to evaluate the safety and efficacy of *CISH* inactivation. Remarkably, the trial reported an ongoing and durable complete response lasting over two years in a young adult patient with colorectal cancer that was refractory to multiple lines of chemotherapy and immunotherapy. This outcome is particularly noteworthy given the grim prognosis typically associated with metastatic colorectal cancer, where treatment options are extremely limited, and survival outcomes are almost uniformly poor. The durable complete response in such a heavily pretreated patient provides strong clinical validation of the potential of *CISH* as a therapeutic target.

Given this landmark clinical result, together with mounting preclinical evidence demonstrating the broad utility of *CISH* targeting across both hematological malignancies and solid tumors, next-generation small molecule drugging strategies aimed at modulating CISH are now in advanced stages of development.

## METHODS

### Cell lines

H-1975 (CRL-5908) and Phoenix-AMPHO (CRL-3213) cells were obtained from ATCC. H-1975 cells stably expressing mutant KRAS^G12D^ were sourced from Horizon Discovery (HD 120-001). The NALM6 FFLuc/GFP reporter cell line was purchased from Creative Bioscience (CSC-RR0360). All cell lines were cultured under recommended media formulations and growth conditions, maintained at sub-confluent densities, and routinely tested for mycoplasma contamination.

### Generation of CD19-expressing NALM6 and H-1975 cells

NALM6 CD19^KO^ cells were generated using a Synthego CRISPR knockout kit targeting the *CD19* gene. To create NALM6 cell lines with varying levels of CD19 expression, CD19^KO^ cells were transduced with a lentiviral vector encoding PGK-driven expression of truncated CD19 (lacking the intracellular signaling domain). Individual clones exhibiting distinct CD19^Low/Med^ expression were selected and sorted for uniform expression profiles. For H-1975 cells expressing CD19, the parental H-1975 cell line (CRL-5908) was transduced with a lentiviral vector encoding EF1A-driven expression of truncated CD19.

### Isolation and culture of primary T cells from healthy donors

Leukopaks were purchased from the NHS Blood and Transplant Bank (NHSBT) from anonymized healthy individuals and handled and stored in accordance with the Human Tissue Authority UK regulations. PBMCs were isolated by density-gradient centrifugation using lymphocyte separation medium (Corning) and cryopreserved until ready for use. Total CD8^+^ T cells were isolated from unfractionated PBMCs using either the EasySep™ or MojoSort™ Human CD8^+^ T Cell Isolation Kit (Stem Cell Technologies, Biolegend) and following manufacturer’s guidelines. Isolated human CD8^+^ T cells were cultured in X-VIVO-15 Basal Media (Lonza) supplemented with 10% Human AB Serum Heat Inactivated (Sigma), 300IU/ml Recombinant Human IL-2 (unless otherwise stated), 5ng/ml Recombinant Human IL-7, and 5ng/ml Recombinant Human IL-15 (all Peprotech) and 10mM N-Acetyl-L-cysteine (Sigma) (henceforth referred to as complete T cell media) and cultured in a 37°C, 5% CO_2_ culture incubator. Media was replaced every 2-3 days with fresh complete media including cytokines.

### Immune checkpoint edited CD8^+^ T cells using CRISPR/Cas9

CD8^+^T cells were stimulated using TransAct (1:100, Miltenyi Biotec, 130-128-758) in complete T cell media and under normal growth conditions for 48-72 hours prior to electroporation. T cells (1-2 million) were electroporated with Cas9–sgRNA–RNP (60 pmol Cas9: 2-4 ug sgRNA) using Lonza’s P3 Primary Cell Nucleofector® Kit (V4SP-3096) program EH-115, following manufacturer’s protocol. Unless otherwise stated, control-edited T cells were targeted with the *AAVS1* sequence. All sgRNA sequences are listed in Table S1. All guide RNAs were ordered as Alt-R CRISPR-Cas9 sgRNAs from Integrated DNA Technologies (IDT) except for PTPN1 and PTPN2, which were purchased as ready Gene-Knock out kits from EditCo.

### Analysis of Gene Knockout Efficiency on DNA Level

Primers for PCR were designed to amplify a 600-900 basepair region surrounding the sgRNA target site. Following a minimum of 48 hours after electroporation, genomic DNA was extracted from CD8^+^ T cells and target regions were amplified by PCR using the GoTaq G2 PCR mastermix (Promega). Correct and unique amplification of the target regions was verified by agarose gel electrophoresis before purifying PCR products using the QIAquick PCR Purification Kit (Qiagen). For analysis by TIDE, PCR amplicons were Sanger sequenced and paired ab1 files of control versus edited samples were analyzed using Synthego’s ICE tool (https://ice.synthego.com). All primer sequences used are listed in Table S1.

### Production of AAV-mediated TCR-Knock-in and checkpoint-knockout CD8^+^ T cells using CRISPR/Cas9

Isolated CD8^+^ cells were thawed, activated and electroporated as described above, followed by a resting period at room temperature for 15 min to allow for cell recovery. After recovery, cells were resuspended in prewarmed complete T cell media. To achieve targeted KRAS^G12D^-TCR knock-in, rAAV6 was added to CD8^+^ T cells 2-5 hours after electroporation at an MOI of 1×10^6^. Viral rAAV6 particles were produced by Signagen. Electroporated T cells were recovered in complete T cell media at a density of 1×10^6^ cells per ml and allowed to rest for 48 hours before subsequent analysis.

### Flow cytometry analysis of T cell phenotypes

For flow cytometric analyses of the CRISPR-edited T cell phenotypes and cell surface marker expression, cells were harvested from culture plates and washed using FACS Buffer containing PBS with 0.5% Bovine Serum Albumin (Thermo Scientific), followed by staining with fluorophore-conjugated monoclonal antibodies against cell surface markers. All antibodies used for flow cytometric analyses are listed in Table S2. Live/Dead Fixable Dead Cell Stains (Invitrogen) were included in all experiments to exclude dead cells. All samples were acquired on an Cytek Aurora cytometer (Cytek^®^ Biosciences), and data was analyzed using FlowJo 10 software (BD Biosciences). All antibodies used for flow cytometric analyses are listed in Table S2.

### Intracellular cytokine staining

Cells were stimulated for a total of 6 hours with anti-hCD3 (OKT3) or anti-mTCRb (H57) (as stated in figure legend). After one hour of stimulation GolgiStop solution (BD Bioscience) was added (a total of 5 hours block of intracellular protein transport). As a positive control for cytokine production, a pool of T cells was stimulated for 6 hours with 50ng/ml PMA and 1ug/ml Ionomycin (Sigma). T cells were then harvested and washed with FACS Buffer and stained for surface markers followed by fixation and permeabilization using BD Cytofix/Cytoperm Fixation/Permeabilization Solution (ThermoFisher) before proceeding with intracellular cytokine staining. Live/Dead Fixable Dead Cell Stains (Invitrogen) were included in all experiments prior to staining to exclude dead cells.

### Realtime Cytolysis Assay (xCELLigence)

Cytolysis assays were performed using the xCELLigence RTCA SP platform (Agilent), which measures electrical impedance to generate a cell index (CI). Background readings were recorded with media alone prior to seeding. Adherent H-1975 tumor cells were plated in 96-well RTCA View plates at a density optimized to enter the linear growth phase after 14–18 hours of incubation at 37°C and 5% CO₂ in complete growth medium. The following day engineered T cells (with indicated gene edits) were added at specified effector-to-target (E:T) ratios. Assays were run undisturbed for up to 90 hours, with impedance measurements taken every 2–10 minutes.

Data were analyzed using RTCA software and expressed as % cytolysis, calculated as: % Cytolysis = [(CI_target only – CI_with effectors) ÷ CI_target only] × 100. Controls included: Background (media only); Negative control (target cells only); Positive control (target cells treated with 2.5% Triton X-100 for maximum cytolysis).

### CD19-CAR^+^ T production

Preparation of retrovirus: CD19-CAR retrovirus (Yescarta^27, 28^) was generated by transient transfection of Phoenix-AMPHO packaging cells. Briefly, 5 million Phoenix-AMPHO cells were plated per 10-cm dish 24 hours before transfection in DMEM + 10% FBS and transfected with 20ug of CD19-CAR plasmid (Yescarta^27, 28^) using Lipofectamine Plus following manufacturer’s instructions. Viral supernatant was collected after 48 hours and loaded onto RetroNectin-coated (10 ug/ml, Takara Bio) non-TC 24-well plates and centrifuged at 2,000*g* for 2 hours at 32°C.

### Engineering of CISH depleted CD19-CAR^+^ T cells

CD8^+^ cells from healthy donors’ PBMCs were stimulated with TransAct (1:100, Miltenyi Biotec, 130-128-758) for 48 hours before KO/transduction. Stimulated cells were then CRISPR-engineered to knock-out immune checkpoints (CISH). Preformed Cas9–sgRNA–RNP (60 pmol Cas9: 2-4 ug sgRNA) and CD8^+^ cells were gently resuspended in P3 buffer with supplement (Lonza’s P3 Primary Cell Nucleofector® Kit, V4SP-3096) at 2 million cells per 20 μl, and pulsed with code EH-115. After electroporation, the 4D-Nucleofector cuvette was placed at room temperature for 15 min to allow for cell recovery. After recovery, cells were resuspended in prewarmed X-vivo media supplemented with 300IU/ml IL-2, 5 ng/ml IL-7 and 5 ng/ml IL-15. Cells were then transduced with CD19-CAR viral supernatant by transferring them onto the RetroNectin/virus coated plates (see above), and spun at 32°C for 30 mins at 300rpm. Cells were then transferred to 37°C incubator to rest overnight. Cells were assessed for transduction efficiency after 3–4 days by measuring surface expression of CD19-CAR by flow cytometry. If needed, CD19-CAR^+^ CD8^+^ cells were enriched using the MACSprep™ CD19 CAR MicroBead Kit following manufacturer’s instructions (Miltenyi Biotec, 130-127-866).

### CD19-CAR^+^ T cells in-vitro Cytolysis Assay

CD19-CAR^+^ CD8^+^ T cells were co-cultured with pre-plated GFP^+^ FFLuc^+^ NALM6 tumor cells in a 96-well flat bottom plate starting at a 1:2 E:T ratio then with a log_2_ serial dilution in triplicates. After 18hrs, NALM6 viability was measured by luciferase.

### Statistical analyses

Appropriate statistical tests were used to analyze data, as described in each figure legend. Statistical differences between two sample groups, where appropriate, were analyzed by a standard Student’s two-tailed, non-paired, t-test and between three or more sample groups using two-way or three-way ANOVA using GraphPad Prism Software version 10. Significance was preset at *P* < 0.05. P and N values are indicated in the figures where statistical analyses have been carried out.

---

**Supplementary Fig 1.**
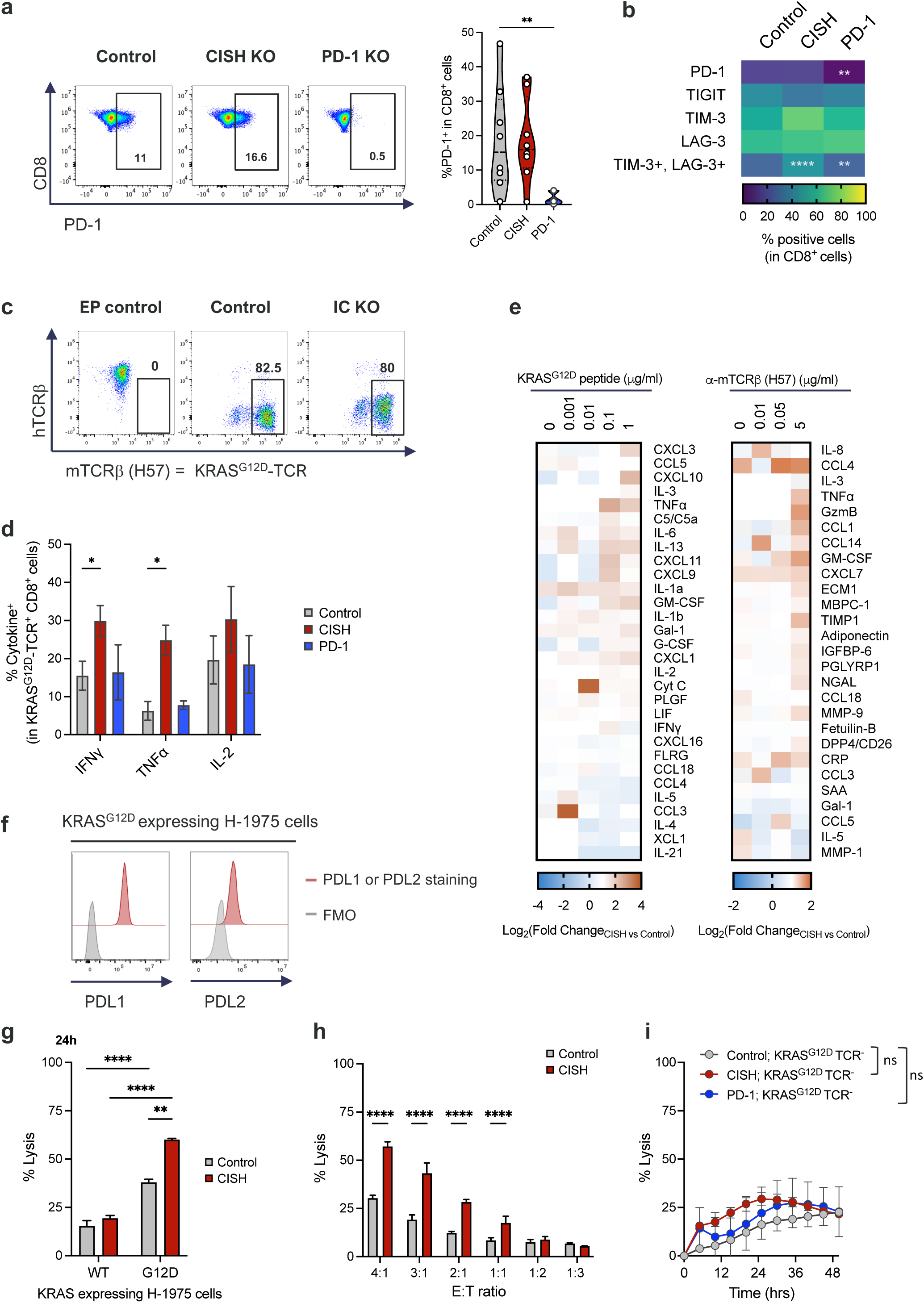
**a**, Validation of PD-1 KO by cell surface flow cytometry staining in CISH and PD-1 depleted CD8^+^ cells. **b,** Heatmap depicting levels of exhaustion markers in unstimulated CISH and PD-1 depleted CD8^+^ cells. **c,** Successful integration of KRAS^G12D^-TCR into *TRAC* locus in immune checkpoint (IC) deleted CD8^+^ T cells after exogenous mTCRβ (H-57) enrichment, as confirmed by flow cytometry. EP: electroporation. **d**, Effector Cytokines: IFN**γ**, TNF**α** and IL-2 production in KRAS^G12D^-TCR^+^ CD8^+^ cells after 24 hrs post stimulation with ⍺-mTCRβ (H57; 1 ug/ml), as measured by intracellular cytokine staining. **e**, Cytokine profiling of CISH depleted KRAS^G12D^-TCR^+^ cells at 24 hours after stimulation with either pulsed KRAS^G12D^ peptide (antigen-specific) or ⍺-mTCRβ (H57) – as measured by nELISA. **f,** Cell surface expression of PDL1/L2 in KRAS^G12D^ expressing H-1975 cancer cells, measured by flow cytometry. **g-i,** Depletion of CISH increases antigen-specific killing in a model of KRAS^G12D^ solid tumor. **g**, Quantification of cytolysis at 24 hours timepoint of H-1975 stably expressing either WT or mutant G12D KRAS, by edited KRAS^G12D^-TCR^+^ CD8^+^ T cells at 4:1 Effector: Target ratio. **h,** Quantification of cytolysis at 6 hours timepoint of KRAS^G12D^ expressing H-1975 cells by edited KRAS^G12D^-TCR^+^ CD8^+^ T cells at various Effector: Target ratios. **i,** Killing of H-1975 target cells stably expressing KRAS^G12D^ by non-transduced (KRAS^G12D^-TCR^-^) CD8^+^ T cells in the presence (Control) or absence of CISH or PD-1, at effector: target ratio 10:1. **a-i:** Statistical significance was determined by either One or Two-way ANOVA vs. Control, with multiple comparison test for repeated measures. Not shown = not significant, *P >0.05, **P>0.01, ***P>0.001, ****P>0.0001. All data are representative of three independent experiments. Data are mean ± SEM.

**Supplementary Fig 2.**
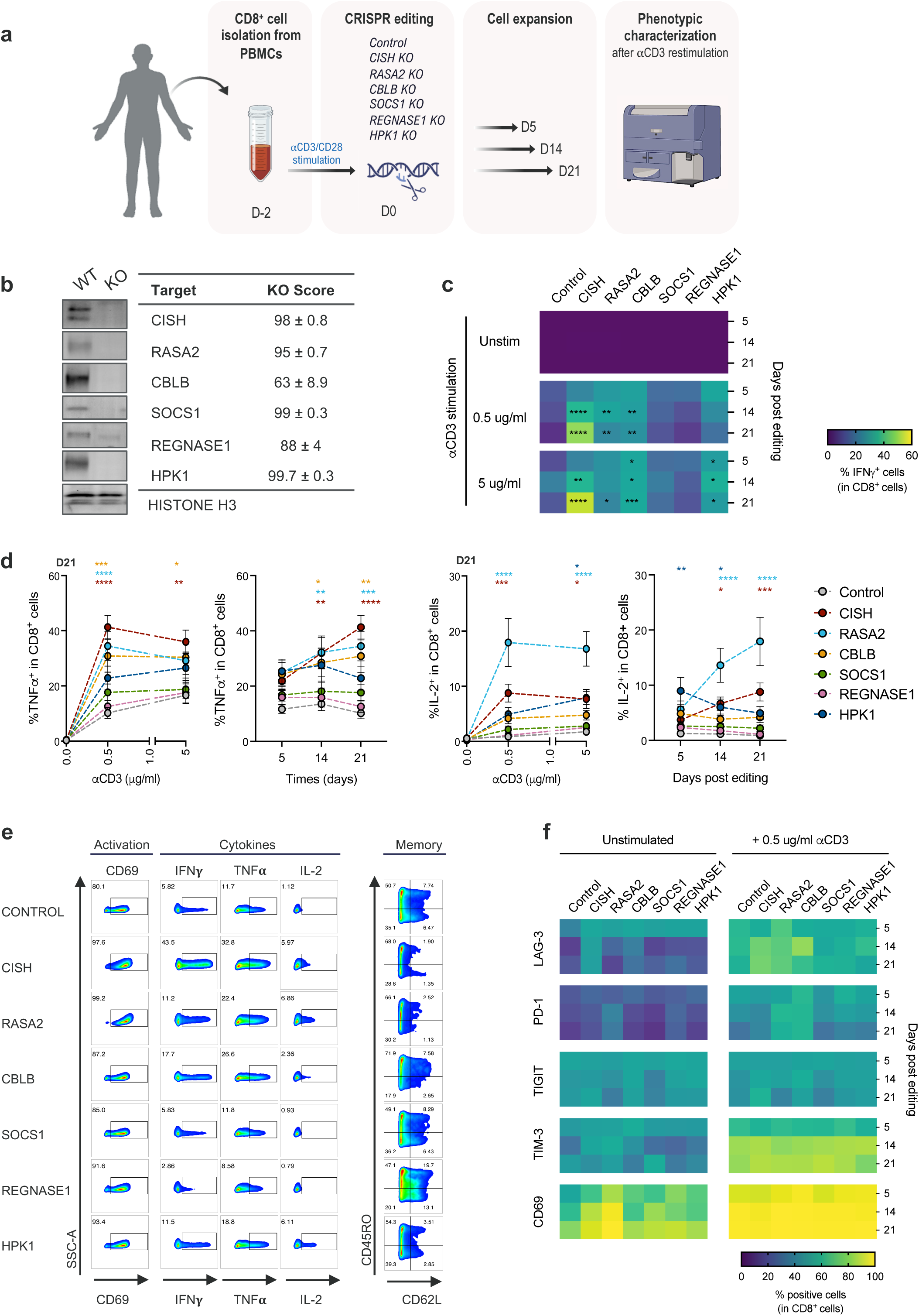
**a,** Schematics of engineering and functional phenotype assessment of primary human CD8^+^ T cells. **b,** Efficient deletion of intrinsic immune checkpoints, as measured at DNA level (KO scores, right panel) and protein level (western blots, left panel). **c,** Heatmap depicting levels of IFN**γ** producing cells after ⍺CD3 restimulation over time post editing. **d,** Evaluation of effector cytokines TNF⍺ and IL-2 production in CD8^+^ cells upon CD3 stimulation (dose response), and over time post editing after 0.5 ug/ml ⍺CD3 stimulation. Statistics shown vs. Control at each ⍺CD3 dose (Right panels) or vs. Control at each time point (Left panels). **e,** Flow Cytometry analysis of activation, cytokine production and memory markers in CD8^+^ cells at day 21 post editing, after 0.5 ug/ml ⍺CD3 stimulation. Graphs are representative of 6 donors. **f,** Heatmap depicting levels of activation and exhaustion markers in unstimulated or 0.5 ug/ml ⍺CD3 stimulated CD8^+^ cells over time post editing. **b-f**: Statistical significance was determined by ANOVA vs Control (unless otherwise stated); *P ≤ 0.05; **P ≤ 0.01; ***P ≤ 0.001; ****P ≤ 0.0001; not shown = not significant. All data is representative of at least three independent experiments, N=6+. Error bars represent mean ± SEM.

**Supplementary Fig 3.**
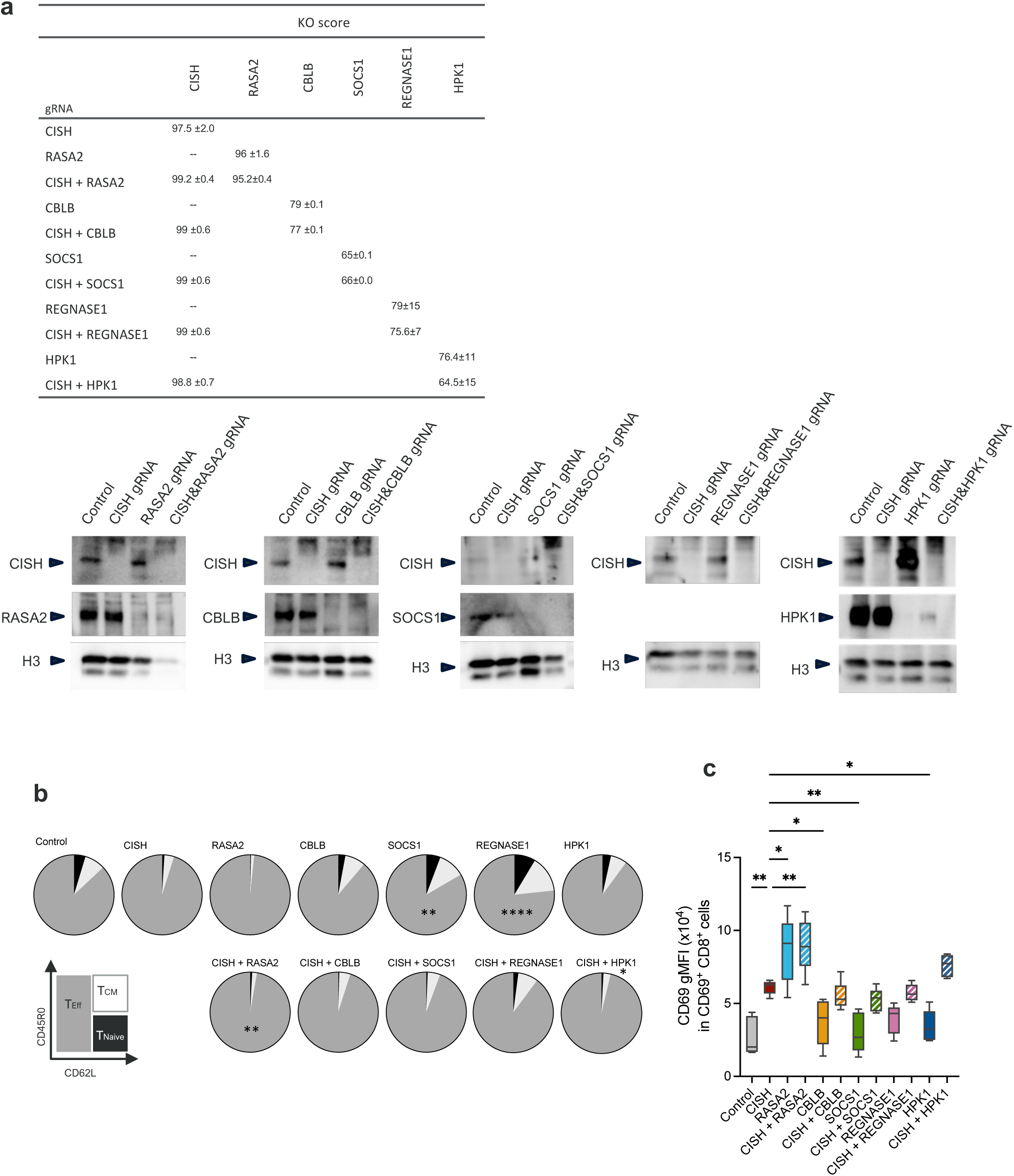
**a,** Efficient deletion of intrinsic immune checkpoints, as measured at protein (western blots, lower panel) and DNA level (table). **b,** Proportion of cells with an effector memory phenotype in CISH depleted CD8^+^ T cells in combination with other intracellular immune checkpoints, at day 21 upon 0.5 ug/ml ⍺CD3 stimulation. **c,** Increased activation status of CISH depleted CD8^+^ T cells in comparison to other intrinsic immune checkpoint deleted CD8^+^ T cells, upon 0.5 ug/ml ⍺CD3 stimulation (Day 21). **b-c**: Statistical significance was determined by One-way ANOVA vs. CISH KO. *P ≤ 0.05; **P ≤ 0.01; ***P ≤ 0.001; ****P ≤ 0.0001; not shown = not significant. All data is representative of at least three independent experiments. Error bars represent mean ± SEM.

**Supplementary Fig 4.**
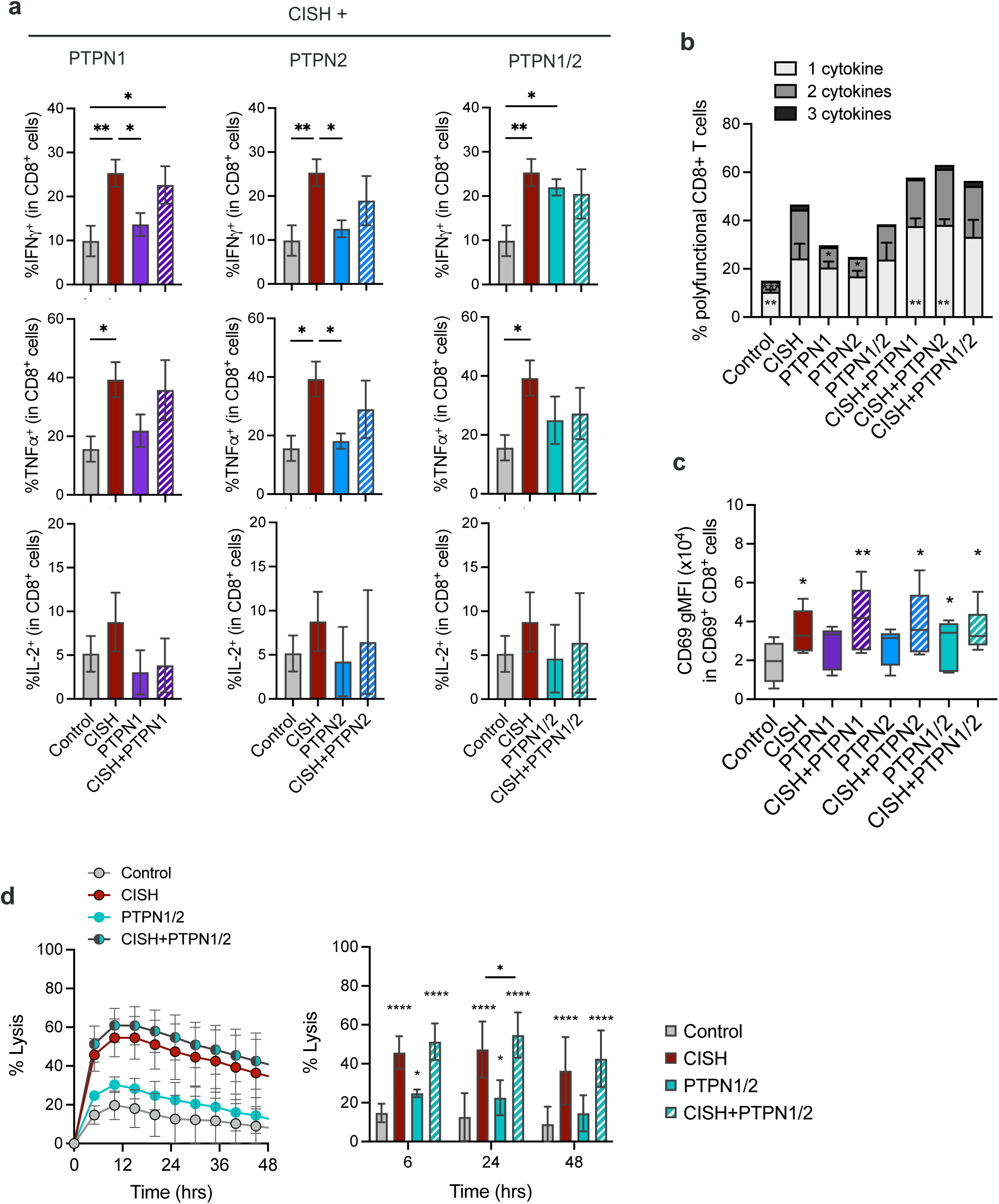
**a**, T cell effector function when CISH is depleted in combination with phosphatases PTPN1/2. Effector cytokines production upon 0.5 ug/ml ⍺CD3 stimulation, measured by ICS. **b**, Polyfunctionality of CD8^+^ T cells after TCR stimulation when CISH is depleted in combination with PTPN1, PTPN1 or PTPN1/2 double KO (0.5 ug/ml ⍺CD3 stimulation). Statistical significance shown vs. CISH KO. **c**, Increased activation status of CISH depleted CD8^+^ T cells in comparison to PTPN1/2 deleted CD8^+^ T cells, upon 0.5 ug/ml ⍺CD3 stimulation. Statistical significance shown vs. Control. **a – c**: Statistical significance was determined by One-way ANOVA. *P ≤ 0.05; **P ≤ 0.01; ***P ≤ 0.001; ****P ≤ 0.0001; not shown = not significant. All data is representative of at least three independent experiments, N=4. Error bars represent mean ± SEM. **d**, Antigen-specific killing in a model of KRAS^G12D^ solid tumor when CISH is depleted in combination with phosphatases PTPN1/2. Real-time measurement of cytolysis of KRAS^G12D^ expressing H-1975 cells by edited KRAS^G12D^-TCR^+^ CD8^+^ T cells. (2:1 Effector: Target ratio). Statistical significance was determined by One-way ANOVA vs. Control, unless otherwise stated in graph. *P ≤ 0.05; **P ≤ 0.01; ***P ≤ 0.001; ****P ≤ 0.0001; not shown = not significant. N=3. Error bars represent mean ± SEM.

**Supplementary Fig 5.**
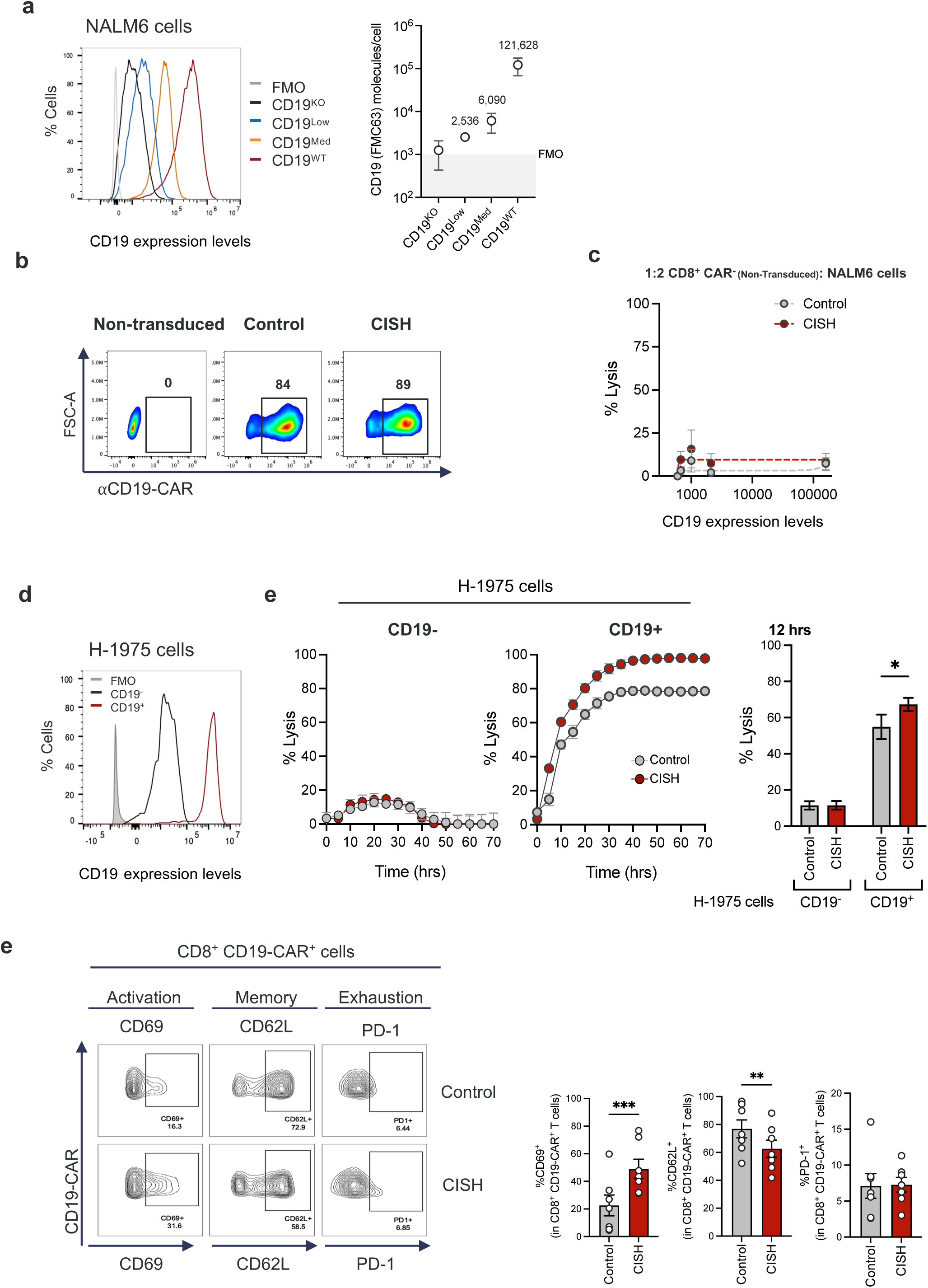
**a,** Generation of CD19 expressing NALM6 cells. Left, Representative histogram plot of CD19 expression of the different NALM6 clones. Right, CD19 density quantification on the surface of NALM6 cells for the different clones. Number of molecules of CD19 were semi-quantitatively determined by the BD Quantibrite Kit. **b**, Flow cytometry data showing levels of CD19-CAR positivity in CD8^+^ cells, after enrichment. Percentages were used to adjust for equal numbers of CAR^+^ T cells per assay. Results are representative of 8 donors. **c,** NALM6 target cell killing by Non-transduced CD8^+^ T cells in the presence (Control) or absence of CISH (CISH) at effector: target ratio 1:2. **d,** Generation of CD19 expressing H-1975 non-small cell lung cancer cell line. Flow cytometry histograms showing CD19 staining in H-1975 cells engineered to express CD19 (in red) compared to unperturbed H-1975 cells (in black). FMO in grey. **e,** Cytotoxicity of CISH depleted CD19-CAR^+^ CD8^+^cells at an Effector: Target ratio of 1:2 using H-1975 cells either unperturbed (CD19^-^) or expressing CD19 (CD19^+^) as target cells. Killing over time using an xCELLigence assay. Representative data from one CD19-CAR^+^ T cell donor for killing of target CD19^+^ H-1975 cells (each time point in triplicates, data are mean ± SD). Data are representative from 2 donors. **f,** Flow Cytometry analysis and quantification of activation, memory and exhaustion markers in CD19-CAR^+^ CD8^+^ cells (day 7-12 post transduction). Results are representative of 8 donors.

**Table S1.**
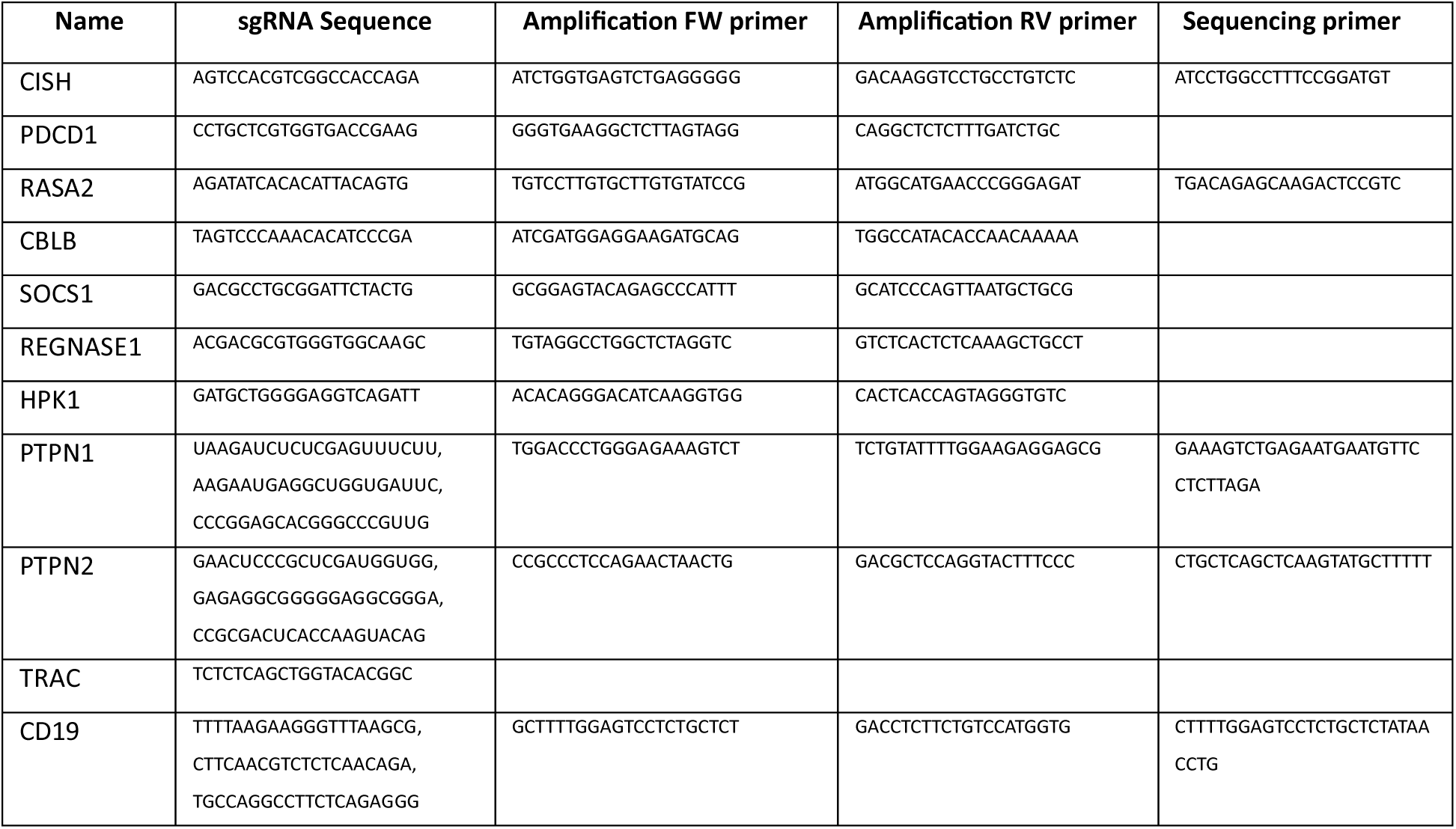
Guide RNA and primer sequences for CRISPR-knockout engineering.

**Table S2.** Antibodies used for flow cytometric analyses.

| Name | Clone<br>(if applicable) | Fluorophore | Vendor | Cat No. | Dilution used |
| --- | --- | --- | --- | --- | --- |
| CD3 | CUCHT1 | FITC | ThermoFisher |  | 1:100 |
| CD4 | OKT4 | BV785 | Biolegend | 317442 | 1:100 |
| CD8a | HIT8a | BV605 | Biolegend | 300936 | 1:100 |
| CD45R0 | UCHL1 | PE-Dazzle594 | Biolegend | 304248 | 1:100 |
| CD45RA | HI100 | APC | Biolegend | 304112 | 1:80 |
| CD62L | DREG-56 | PE-Cy7 | Biolegend | 304822 | 1:50 |
| CD69 | FN50 | APC | Biolegend | 310910 | 1:50 |
| 4-1BB | 4B4-1 | AOC | Biolegend | 309810 | 1:50 |
| Granzyme B | QA16AO2 | APC/Fire™ 750 | Biolegend | 372210 | 1:50 |
| LAG-3 | 11C3C65 | BV785 | Biolegend | 369309 | 1:80 |
| TIGIT | A15153G | PE/Dazzle 594 | Biolegend | 372716 | 1:50 |
| PD-1 (CD279) | EH12.1 | BV711 | Biolegend | 329928 | 1:50 |
| IFN-γ | 4S.B3 | FITC or AF647 | Biolegend | 502506/502516 | 1:50 |
| TIM3 | F38-2E2 | PE or BV711 | Biolegend | 345006/345024 | 1:80 |
| TNF-α | MAB11 | PE or AF488 | Biolegend | 502909/502915 | 1:50 |
| IL-2 | MQ1-17H12 | BV421 | Biolegend | 500328 | 1:50 |
| hTCRa/b | IP26 | APC | Biolegend | 306718 | 1:50 |
| mTCRb | H57-597 | PE | Biolegend | 109208 | 1:100 |
| CD19 | FMC63 | PE | Novus Bio | NBP2-52716PE | 1:100 |
| CD19-CAR FMC63 | REA1297 | PE | Miltenyi Biotec | 130-127-342 | 1:100 |

## Notes

### Competing Interest Statement

The authors have declared no competing interest.

